# Discovering cell types underlying rare disease phenotypes using scRNA-seq data from non-diseased tissues

**DOI:** 10.64898/2025.12.09.693155

**Authors:** Jorge Novoa, Florencio Pazos, Monica Chagoyen

## Abstract

Despite their low individual prevalence, rare diseases collectively pose a significant health burden, affecting millions of people worldwide. These conditions often result from single-gene mutations, yet the cellular contexts in which these alterations act remain largely unknown—information crucial for improving diagnosis and treatment. As patient-derived samples are scarce, we use single-cell RNA sequencing (scRNA-seq) data from non-diseased tissues to identify relevant cell populations. We introduce Cell4Rare, a computational framework that integrates these healthy scRNA-seq datasets with known phenotype-associated genes to map disease phenotypes to specific cell types. Applied across diverse tissues and phenotypes, Cell4Rare was validated against literature-based associations, achieving robust performance with an AUC of 0.71. These results highlight the potential of computational analyses of non-diseased scRNA-seq data to uncover the cellular basis of rare disease phenotypes, paving the way for improved diagnostics and therapeutic strategies.

## 1. Introduction

Rare diseases, despite their individual low prevalence, collectively affect millions worldwide and pose persistent challenges for both diagnosis and treatment. The majority of these conditions have a genetic basis, with mutations that disrupt molecular and cellular pathways, ultimately manifesting as distinct and often severe phenotypes. Experimental approaches such as CRISPR-Cas9 gene editing, animal models, patient-derived iPSCs, and in vitro functional assays have become indispensable tools for validating the pathogenicity of specific variants and establishing the molecular and cellular pathways disrupted (MacArthur et al., 2014). These strategies provide critical insights into disease etiology by enabling the direct interrogation of gene function and phenotype. However, they are often limited by high costs, extended timelines, and scalability challenges—particularly in the context of rare diseases, where patient samples and resources are scarce. These limitations underscore the urgent need for robust, scalable computational tools that make use of the more abundant healthy omics data to facilitate the discovery and understanding of rare disease–causing mechanisms.

A major obstacle in rare disease research is determining the specific cellular contexts in which pathogenic genetic variants exert their effects. This is especially challenging given that many rare disease–associated genes are broadly expressed across tissues but lead to phenotypes that are restricted to a small subset of cell types (Feiglin et al., 2017). Pinpointing these relevant cellular targets is crucial for understanding disease mechanisms and for designing effective, targeted therapies. Recent advances in single-cell transcriptomics, along with growing databases of genotype–phenotype associations, offer an unprecedented opportunity to study gene activity at cellular resolution. Yet, for most rare diseases—many of which have pediatric onset—single-cell data from affected individuals remain unavailable, making traditional case–control approaches impractical.

In this context, computational methods that extract mechanistic insights from single-cell data derived from non-diseased tissues offer a valuable alternative. These approaches can help infer the cellular consequences of genetic alterations and guide the selection of the most relevant experimental systems for downstream validation. Prior studies have successfully used single-cell RNA sequencing (scRNA-seq) from healthy tissues to identify disease-relevant cell types in common disorders such as schizophrenia (Skene et al., 2018), Parkinson’s disease (Bryois et al., 2020), and also several rare genetic conditions (Hekselman et al., 2024). Building on this foundation, our group previously introduced a hypothesis-driven framework that prioritizes disease-relevant cell types based on gene co-expression and phenotype-associated gene sets (Alías-Segura et al., 2024). We found that co-expression analyses outperformed differential expression in recovering known gene–phenotype associations. However, this approach required constructing co-expression networks for each cell cluster—a time-consuming and computationally intensive process that limited its scalability. Moreover, the resolution was limited to the level of cell types or clusters, rather than individual cells.

To address these limitations, we introduce Cell4Rare, a novel computational framework that systematically maps the effects of genetic alterations to specific cell types implicated in disease phenotypes. By integrating single-cell transcriptomic data with genetic etiologies and clinical phenotype information, Cell4Rare identifies the cell populations most likely to be impacted by disease-causing mutations. It does so by modeling gene expression in a cell type–specific manner, capturing the context-dependent impact of genetic variants and offering mechanistic hypotheses that link genotype to phenotype.

We applied Cell4Rare to a large compendium of genetic disorders—including many rare diseases—with well-characterized gene–phenotype relationships. Our results reveal a strong correspondence between predicted affected cell types and known disease pathophysiology. Importantly, Cell4Rare can resolve effects down to the level of individual cells, generating novel, testable hypotheses for experimental follow-up and therapeutic targeting.

By systematically linking genetic variation to cell type–specific functional effects, Cell4Rare addresses a central challenge in rare disease research. It offers a scalable, data-driven approach to uncover the cellular basis of genotype–phenotype relationships, with broad implications for improving diagnostic precision and guiding the development of targeted therapies. In this study, we describe the methodological foundations of Cell4Rare, validate its predictions across diverse datasets, and explore its potential to advance our understanding of rare disease biology.

## 2. Results

The Cell4Rare approach builds on our previous work (Alías-Segura et al., 2024), where we showed that gene co-expression patterns better capture known gene–phenotype associations than standard differential expression analyses. In this study, we extend that principle by investigating whether individual cells of a given cell type tend to express a greater number of phenotype-associated genes compared to other cells within the same tissue.

Since the number of genes from any gene set detected in a cell is inherently influenced by the cell’s overall transcriptional activity, we first model the expected relationship between the total number of expressed genes and the number of genes associated to a given phenotype using linear regression. Each cell is then evaluated based on its deviation from this expected trend, allowing us to quantify whether it expresses more phenotype-relevant genes than anticipated given its global expression profile. To detect statistically significant enrichment patterns across cell populations, we applied a signed two-sample Kolmogorov–Smirnov (KS) test, comparing the distribution of deviations within each cell type or cluster to that of all other cells. This enables the identification of cell types or subtypes that consistently express phenotype-associated genes.

To evaluate the Cell4Rare method, we applied it to single-cell RNA sequencing (scRNA-seq) data from non-diseased human tissues compiled in the Human Protein Atlas (HPA) (Digre & Lindskog, 2023) and Tabula Sapiens (TS) datasets (Jones et al., 2022). We used the provided cell-type annotations for both datasets and, in the case of HPA, also incorporated cell cluster definitions to capture finer-grained transcriptional heterogeneity. For each tissue, we curated a set of relevant phenotypes from the Human Phenotype Ontology (HPO) (Köhler et al., 2020), along with their associated gene sets. Phenotypes were selected based on anatomical mappings provided by the HPO and further refined through manual curation, as described in the Methods section and Supplementary Material.

We then applied Cell4Rare to identify cell types and clusters showing statistically significant enrichment for phenotype-related gene expression. Multiple testing correction was applied to control the false discovery rate. Finally, we validated our predictions by comparing them to known phenotype–cell type associations automatically compiled from the literature. An overview of the analytical workflow is provided in Figure 1 (see also the Methods section for additional details).

**Figure 1.**
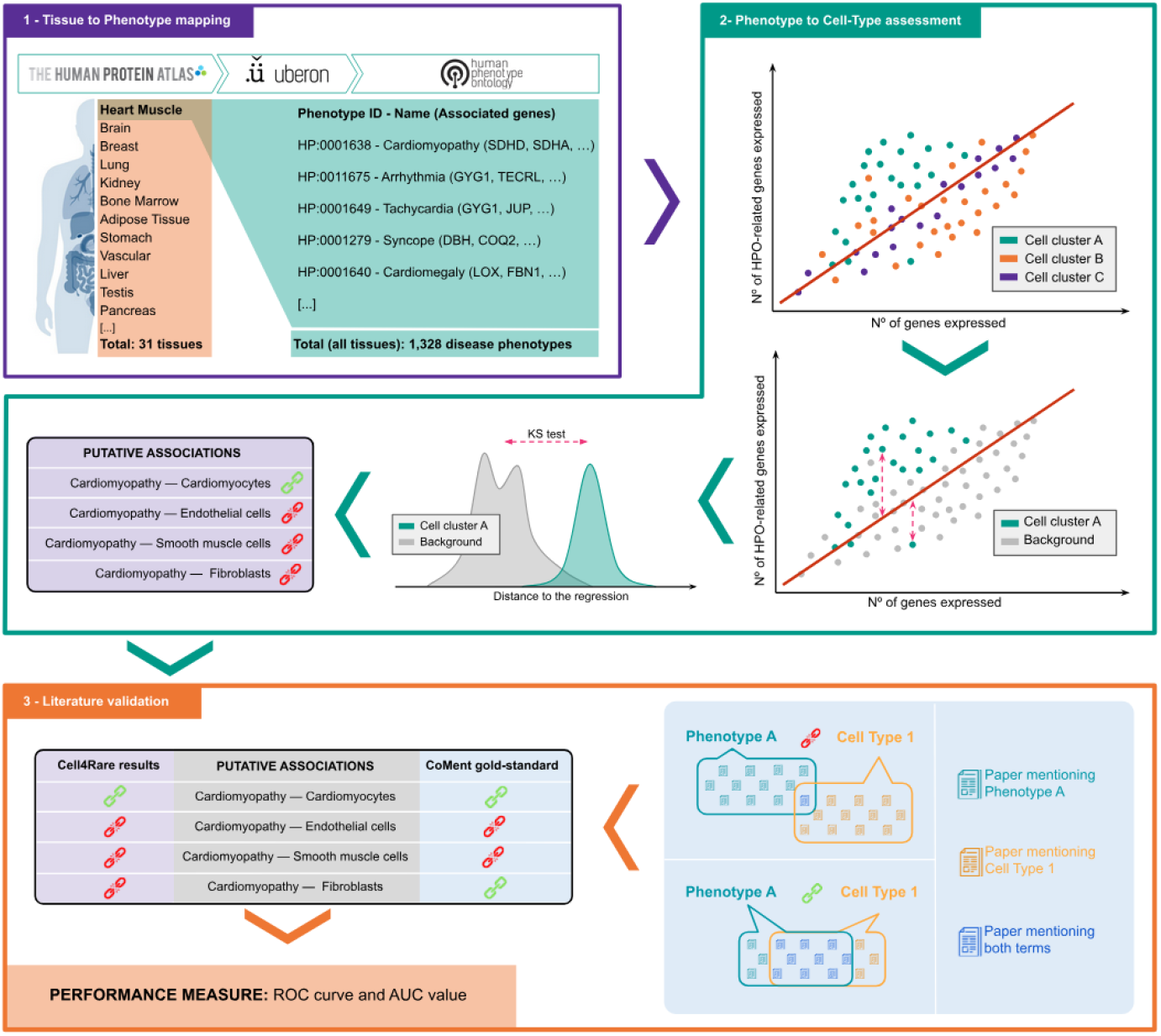
Overview of the methodology. (1) Tissue to phenotype mapping. We obtained scRNA-seq data of 31 non-diseased human tissues from the HPA and mapped each tissue to its associated phenotypes. **(2) Phenotype to cell-type assessment**. For each tissue-phenotype pair, we evaluated whether the phenotype is associated with a given cell type or cell cluster. **(3) Literature validation**. We validated the resulting phenotype-cell type associations using CoMent.

Here, we present the results derived from the HPA dataset. Analyses based on the TS dataset are provided in the Supplementary Material.

### 2.1. Inferred Phenotype–Cell Type and Cluster Associations

We analyzed single-cell RNA-seq data spanning 31 human tissues curated by the HPA (Digre & Lindskog, 2023), aggregated from multiple independent studies (Bhat-Nakshatri et al., 2021; Chen et al., 2018; De Micheli et al., 2020; Guo et al., 2018; he et al., 2020; Hildreth et al., 2021; Jones et al., 2022; Liao et al., 2020; Lukassen et al., 2020; MacParland et al., 2018; Menon et al., 2019; Parikh et al., 2019; Qadir et al., 2020; Solé-Boldo et al., 2020; Ulrich et al., 2022; Vento-Tormo et al., 2018; Wagner et al., 2020; L. Wang et al., 2020; W. Wang et al., 2020; Y. Wang et al., 2019). The HPA dataset includes annotations for 557 distinct cell clusters, corresponding to 85 unique cell types. We mapped a total of 2,630 phenotypes, including 1,328 non-redundant phenotypes — defined as the most specific HPO terms meeting the gene count threshold described in the Methods. Some phenotypes were mapped to multiple tissues. Details regarding the number of phenotypes analyzed per tissue and the phenotype size distribution are provided in Supplementary Figure S3.

Using the HPA dataset, we evaluated 27,603 phenotype-cell type pairs (Supplementary Table S1), identifying 10,706 statistically significant positive associations at a false discovery rate (FDR) < 0.001. These correspond to 9,382 unique phenotype–cell type associations. Results from the TS dataset are presented in Supplementary Table S2 and discussed in the Supplementary Material.

As a representative example, our analysis of heart muscle scRNA-seq data revealed a strong and specific association between *dilated cardiomyopathy* (HP:0001644) and cardiomyocytes (Figure 2a). No significant associations were observed with other cardiac cell-types present in the dataset, including endothelial cells, fibroblasts, and smooth muscle cells. Dilated cardiomyopathy is characterized by ventricular dilation and systolic dysfunction, often progressing to heart failure. This condition is known to have diverse genetic etiologies, many of which impair cardiomyocyte structure and contractility (Favalli et al., 2016).

**Figure 2.**
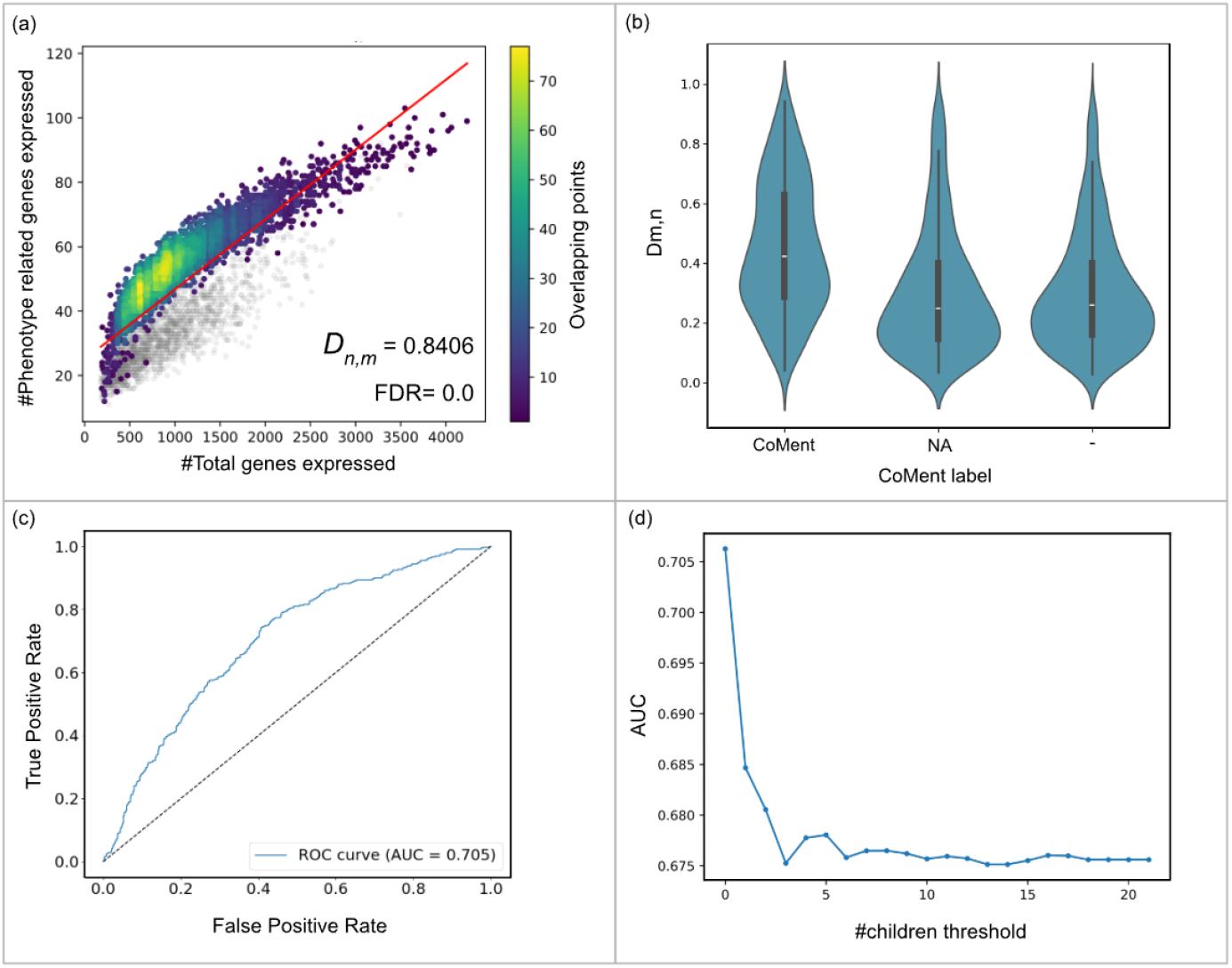
Overview of Cell4Rare results on HPA data. (a) Cell4Rare results for dilated cardiomyopathy analysed using HPA heart muscle scRNA-seq data. Colored points represent cardiomyocytes, and grey points represent other cell types. (b) Distribution of the maximal D_n,m_ value per cell type grouped by literature co-mention. “CoMent” indicates significant co-mention, “–” indicates non-significant or absent co-mention, and “NA” indicates no available information. (c) ROC curve and AUC value for the CoMent-based literature validation of Cell4Rare results on HPA data, following the “Best cell type per phenotype” criterion. (d) AUC values when upper-level phenotypes are included in the analysis.

In the HPA dataset, certain cell types are represented by multiple cell clusters. These cases of multiple clusters may reflect transcriptionally distinct subtypes within a given cell type (e.g., different types of t-cells in PBMC and other tissues or functionally distinct neurons in the brain). To explore this additional layer of heterogeneity, we extended our analysis to infer associations between phenotypes and individual cell clusters. In total, we evaluated 86,756 phenotype–cell cluster pairs (see Supplementary Table S3), identifying 32,694 statistically significant positive associations at FDR < 0.001.

Of particular interest are those cases where a cell type is not significant for a phenotype, but a particular cell cluster within the cell type is. This is the case for 6,941cell-clusters (See Supplementary Table S5). Some of the most extreme examples include specific clusters of bronchus ciliated cells and bronchus club cells related to different respiratory pathologies like recurrent pneumonia, bronchiectasis and cough. Also, specific clusters of brain excitatory neurons related to psychiatric problems including abnormal prosody, aplasia of the olfactory bulb and dementia. As shown in Supplementary Figure S7, the highest *D*_*n,m*_ value for a phenotype more frequently corresponds to a cell-cluster than a cell-type.

To quantify phenotype–cell set associations, we used the test statistic *D*_*n,m*_ from the signed two-sample Kolmogorov–Smirnov (KS) test as a prediction score (see Methods). This statistic captures the divergence between two empirical distributions: one representing gene expression in a given cell population (cell type or cluster), and the other representing all remaining cells in the tissue. Because we applied a signed version of the test, higher values of *D*_*n,m*_ reflect a greater number of phenotype-associated genes being expressed in the cell population of interest.

To evaluate potential biases in our results, we assessed the correlation between phenotype size—defined as the number of genes associated with each phenotype—and *D*_*n,m*_. The Pearson correlation coefficient was –0.016 (P = 5.43 × 10^−9^), indicating a negligible association despite the statistical significance due to large sample size. Similarly, the correlation between the number of cells and *D*_*n,m*_ yielded a Pearson coefficient of –0.091 (P = 1.44 × 10^−232^). These results suggest that *D*_*n,m*_ is not meaningfully biased by either the number of associated genes or the number of cells.

### 2.2. Literature validation

To evaluate the coherence of our results with the current knowledge about disease phenotypes, we compiled a set of known relationships between phenotypes and cell-types inferred from the literature using CoMent (Novoa et al., 2024). We considered phenotype-cell types pairs significantly co-mentioned in the literature (CoMent P-value < 0.001) as positive relations and all remaining pairs as negative relations. Bias was observed when comparing the size distribution of non-redundant phenotypes with at least one cell type association in the literature (used for validation) to those with none (P-value = 1.906 × 10^−6^). For redundant phenotypes and for the totality of the phenotypes no significant bias was observed (P-values = 0.583; P-value = 0.058 respectively).

We compared the positive relations provided by CoMent to those inferred by our method generating a *Receiver Operator Curve* (ROC) and quantified the performance of the method by calculating the *Area Under the Curve* (AUC). For this validation, we established three independent criteria to select these best score: (a) the cell-cluster with the highest *D*_*n,m*_ value among cell-clusters; (b) the cell-type with the highest *D*_*n,m*_ value among cell-types and, (c) either the cell-type or the cell-cluster with the highest *D*_*n,m*_ value among cell-clusters and cell-types. We considered as positive only one relation per phenotype, independently of the tissue (i.e. for phenotypes mapped to multiple tissues, only the highest *D*_*n,m*_ value was considered). The distribution of *D*_*n,m*_ values for each criteria is shown in Figure 2b. The best performing criterion was the Best Cell-Type criterion, with an AUC=0.71 (Figure 2c). The distribution of *D*_*n,m*_ values and ROC curves of the Best Cell-Cluster criterion (AUC=0.66) and the Best Cell-Cluster or Cell-Type criterion (AUC=0.69), along with other explored criteria, are presented in Supplementary Figures S5 and S6.

As shown in Figure 2d, the best performance is achieved when only non-redundant phenotypes were included (most specific). Allowing more general phenotypes (with higher numbers of children within those analyzed) results in decreasing AUC values.

We checked if the lexical complexity of a phenotype might bias its chance of being mentioned in the literature. We found significant negative correlation between phenotypes mentioned at least once in the literature and their lexical complexity, whether by their number of words (signed KS test *D*_*n,m*_ = 0.627, p = 0.0) or by their number of characters (signed KS test *D*_*n,m*_ = 0.59, p = 0.0).

It’s worth noticing that positive relations revealed by our method and not significantly commentioned may be still described in the literature. This is the case, for example, for the phenotype *Productive cough*, that we found to be associated with ciliary cells in lung and bronchus and, although they are not significantly commentioned, their association is widely found in the literature. As described by Hill et al. (2022), malfunctions of the cilia impede normal mucus clearance on the airways which often lead to chronic productive cough. We also found putative association not supported by commenting between *Nevus flammeus* and endothelial cells. Nevus flammeus are congenital red stains on the skin caused by an abnormal dilation and malformation of the skin capillary (Lee & Chung, 2018), therefore its association with endothelial cells is plausible.

## 3. Discussion

In this study, we investigated how gene expression profiles at single-cell resolution can inform the identification of cell types affected in diseases that share a specific phenotypic manifestation. We focused on genetic disorders, many of which are classified as rare diseases, with known pathogenic variants that exert strong effects on phenotype. To this end, we analyzed transcriptional activity in non-diseased human tissues. This choice was motivated by two key considerations: first, non-diseased tissue data are broadly available, which is especially critical for the study of rare diseases, where diseased tissues are often inaccessible; and second, we hypothesize that similar phenotypic outcomes—regardless of the underlying genetic cause—should emerge from disruptions in shared, cell-type–specific functional programs. Moreover, analyzing phenotypes associated with diseases of known genetic etiology offers an opportunity to uncover the cellular basis of similar phenotypic features observed in more common, multifactorial conditions, where the interplay between genetic and environmental influences remains poorly understood.

As in our previous work (Alías-Segura et al., 2024), the primary goal of this study was to identify the specific cell types within a given tissue that are most relevant to the manifestation of a phenotype, while intentionally excluding other tissues. However, certain phenotypes were analyzed in multiple tissues according to their classification in the HPO. The *D*_*n,m*_ statistic allows estimating the extent to which constituent cell types preferentially express phenotype-associated genes. For example, *Facial diplegia*—characterized by bilateral paralysis or weakness of the facial muscles—is classified by the HPO as both a nervous system and musculoskeletal abnormality. According to our analysis, the genes associated with this phenotype are strongly associated with skeletal myocytes in skeletal muscle tissue (*D*_*n,m*_ =0.94) compared to other skeletal muscle cell types. In contrast, they are only moderately associated with inhibitory neurons (cluster 30), the best scoring cluster in the brain (*D*_*n,m*_ =0.26). Similarly, *Anorexia*, mapped to both the nervous and digestive systems, showed strong associations with immune cells in digestive tract tissues (colon, stomach, rectum, and small intestine), with *D*_*n,m*_ values ranging from 0.40 to 0.70, and weak association with microglial cells —the best match identified in the brain—at *D*_*n,m*_ =0.28.

While these results may suggest a stronger link to particular cell types in one system, they do not exclude the possibility that phenotypes such as these may result from disruptions in multiple systems, either simultaneously or independently. *Difficulty walking* provides an illustrative example. Among the two tissues to which this phenotype is mapped in the HPO, the highest Cell4Rare score was observed in skeletal myocytes from skeletal muscle tissue (*D*_*n,m*_ =0.87, with 133 phenotype-associated genes expressed), followed by neuronal cells in brain tissue (*D*_*n,m*_ =0.51, with 223 associated genes). The larger set of phenotype-associated genes expressed in neuronal cells—many of which are not detected in skeletal muscle—suggests the involvement of distinct mechanisms. This pattern may reflect the existence of at least two mechanistically different classes of disorders with this phenotype: one driven by muscular dysfunction, and another by neuronal impairment.

Our study has several limitations arising from both the underlying data and the validation strategy employed that affects the results obtained and the performance reported for Cell4Rare. Gene–phenotype associations in the HPO are primarily derived from gene–disease links, which may not always reflect direct causality. In some cases, the annotated phenotype may be a secondary consequence of the disease rather than a direct effect of the gene itself. However, such distinctions are not captured in the HPO. Despite this limitation, we believe that our statistical framework is sufficiently robust to tolerate this level of noise and still extract meaningful associations.

The scRNA-seq datasets analyzed here represent adult tissues and may not capture gene relationships relevant to all the phenotypes under investigation. Low Cell4Rare scores can nonetheless provide valuable insights into the relevance of specific tissues for the phenotypes under investigation. To illustrate this, we highlight several phenotypes whose maximal *D*_*n,m*_ values fall within the bottom 10%. Such results suggest that none of the cell types represented in the dataset are specifically relevant to the manifestation of the phenotype, i.e. cell populations responsible for the phenotype are not adequately captured. *Frontal bossing* (a prominent or bulging forehead), analyzed in brain tissue, may instead reflect developmental processes or skeletal abnormalities to be better studied in embryonic tissues. Phenotypes such as *synophrys* (unibrow), *highly arched eyebrows*, and *low anterior hairline*, analyzed in skin tissue, may depend on cell types or structures (e.g., hair follicles) that are poorly represented in the adult tissue samples analyzed. *Myelodysplasia* and *acute myeloid leukemia*, although analyzed in bone marrow, may involve rare or dynamically shifting hematopoietic populations not well captured in healthy adult datasets. Complex psychiatric and behavioral phenotypes like *loss of speech, suicidal ideation, dyscalculia* or *tic* may be better explain by specific functional groups of neurons that are not reflected by the very general annotation of the data (only *inhibitory neurons* and *excitatory neurons*).

The quality of cell-type annotation strongly influences the outcome of our method. This is particularly evident for six HPA tissues—lung, prostate, salivary gland, thymus, tongue, and vascular—that are derived from Tabula Sapiens (TS). In these cases, the underlying scRNA-seq data are identical, but the cell annotations differ. As shown in Supplementary Figure S8, the top *D*_*n,m*_ values for each phenotype mapped to these tissues vary substantially depending on which dataset is used.

Even in tissues present in both datasets but with different underlying scRNA-seq data, the impact of cell representation and annotation remains apparent. For example, the TS skin dataset lacks cells annotated as keratinocytes, containing only a small fraction (1.1%) of epithelial cells. In contrast, the HPA skin dataset includes suprabasal and basal keratinocytes, which together account for 15% of the cells in the tissue. Since keratinocytes play a key role in skin physiology, analyses of skin phenotypes using TS data are expected to yield less reliable results, simply due to the limitations in cell-type representation in the dataset.

On the validation, one major challenge lies in the mapping of ontology terms—both for assigning cell types and for linking phenotypes to relevant tissues. Some cell types in the HPA dataset could not be reliably mapped to Cell Ontology (CL) terms, including categories such as *mixed immune cells, undifferentiated cells*, and *mixed cells*. While some of these ambiguously labeled cell types, such as mixed immune cells, may be biologically relevant—as suggested by high scores for *Adenocarcinoma of the colon* (0.65) and *Recurrent infection of the gastrointestinal tract* (0.62)—they are necessarily treated as false positives during validation due to the lack of precise ontology alignment. Similarly, mappings from HPO phenotypes to tissues can occasionally produce implausible associations—for example, *male infertility* being mapped to the fallopian tube and endometrium, which yielded high scores for ciliated cells.

The set of phenotype-cell type associations extracted from the literature using the CoMent approach is also subject to limitations. Polysemous terms can lead to erroneous matches (e.g. *Irritability*, as an emotional state, can be matched with papers related to physical irritability, like *Bowel irritability*), and genuine associations may be missed if they are mentioned in only a small number of publications (as illustrated by the examples commented in Results). Despite these issues, CoMent remains, to our knowledge, the most comprehensive available resource for systematically linking disease phenotypes to specific cell types in the absence of curated gold-standard datasets.

Our study demonstrates the feasibility of computationally identifying cell types implicated in genetic disorders using non-disease tissues and associated patient phenotypes. This approach is particularly valuable for rare disease research, where access to patient-derived samples is often limited. These computational predictions can guide the development of biologically relevant *in vivo* or *in vitro* models, enabling validation of genetic causes and mechanistic investigation of disease pathology. Furthermore, our findings enable more precise diagnostic strategies by focusing on cell type–specific markers rather than relying on whole-tissue analyses or nonspecific sampling. Such precision could also inform the design of targeted therapeutic interventions, selectively modulating the most critical cell populations. Collectively, these advances will facilitate the development of more effective, personalized healthcare solutions for individuals affected by rare diseases.

## 4. Methods

### 4.1. Single-cell expression data

We obtained single-cell transcriptomic data (scRNA-seq) from non-diseased human tissues from two sources: (a) 31 tissues from the Human Protein Atlas (HPA) (https://v23.proteinatlas.org/; Digre & Lindskog, 2023; version 23.0;) and (b) 24 tissues from Tabula Sapiens (TS) (https://tabula-sapiens.sf.czbiohub.org/; Jones et al., 2022; 2022 version). For each tissue, we also retrieved the corresponding cluster and cell-type annotations provided by the respective datasets.

Disease phenotypes and their associated gene annotations were obtained from the Human Phenotype Ontology (HPO) (https://hpo.jax.org/, Köhler et al., 2020; v2024-01-16). Anatomical entities were retrieved from the Uberon Multi-species Anatomy ontology (UBERON) (https://www.ebi.ac.uk/ols4/ontologies/uberon; Mungall et al., 2012; v2024-01-27).

### 4.2 Tissue to phenotype mapping

To establish a compendium of disease phenotypes associated with each tissue, we used two complementary approaches: (a) automated mapping based on UBERON relations and (b) manual curation.

For the UBERON-based mapping (a), we first annotated each tissue with its corresponding UBERON term and expanded this annotation to include all descendant terms present in the UBERON.*obo* file. We then retrieved all HPO terms associated with each UBERON term using the HPO.*owl* file. For each of these HPO terms, we additionally obtained all descendant phenotypes from the HPO.*obo* file. Criteria used to systematically perform the manual annotation (b) are provided in Supplementary Table S6. The final tissue–HPO mappings were constructed by combining the outputs of both approaches. This procedure was performed independently for HPA and TS tissues.

For each selected phenotype, we retrieved the associated gene sets from the HPO database, which aggregates gene–disease associations derived from OMIM (Amberger et al., 2014) and Orphanet (Rath et al., 2012). Specifically, we obtained the list of genes related to each phenotype from the HPO annotation file *phenotype_to_genes*.*txt*. For each tissue–phenotype pair, we computed the maximum number of phenotype-associated genes expressed by any individual cell in the corresponding scRNA-seq dataset, considering a gene as expressed if its transcript count exceeded zero. Phenotypes for which this maximum number of expressed genes was fewer than ten were excluded, as such sparse representation would likely yield noisy results in subsequent analyses.

Finally, for each tissue, we selected only the most specific, non-redundant phenotypes from the resulting list—i.e., those phenotypes that have no children among the set of phenotype terms associated with that tissue. This constituted the final set of phenotypes used in downstream analyses.

### 4.3 Expression analysis

We used the scRNA-seq transcript counts to determine which cell groups (clusters and cell types) within a tissue expressed more genes related to a given phenotype than expected based on their total number of expressed genes. For each phenotype associated with a tissue, we constructed a two-dimensional space in which each cell was represented by two coordinates: (x) the total number of genes expressed by the cell, and (y) the number of phenotype-related genes expressed by the cell. A gene was considered expressed if its transcript count was greater than zero.

We then performed a linear regression on this set of points. Cells located above the regression line were interpreted as expressing more phenotype-related genes than expected. For each cell, we computed the difference between the observed number of phenotype-related genes (y) and the expected value predicted by the linear model. This yielded a distribution of distances to the regression line for all cells in the tissue.

### 4.4 Statistical assessment

We performed a signed two sample Kolmogorov-Smirnov test (KS) for each cell group (cluster or cell type) to assess whether the distribution of distances of that group was significantly higher than the distribution of distances for all remaining cells. Resulting P-values were corrected for multiple testing using the Benjamini-Hochberg false discovery rate (FDR). Cell-types or clusters were considered associated with a phenotype if FDR < 0.001.

For the subsequent literature-based evaluation, we defined three independent criteria to identify positive phenotype-cell group associations in our results: (a) selecting only the association between each phenotype and the cell type with highest D_n,m_ value; (b) selecting only the association between each phenotype and the cell cluster with the highest D_n,m_ value and (c) selecting the association between each phenotype and whichever had the higher D_n,m_ value: the top-ranked cell type or the top-ranked cell cluster. As Tabula Sapiens lacks cluster annotations, only criterion (a) was applied to TS data.

### 4.5 Literature analysis

We obtained a set of known phenotype-cell type associations from the literature using CoMent (Novoa et al., 2024), a tool that identifies significant co-mentions between biomedical concepts across abstracts in the entire PubMed corpus. A detailed description of the method is provided in Pazos et al. (2022). To align our results with the associations compiled from CoMent, each cell-type annotation in the scRNA-seq datasets was mapped to a corresponding Cell Ontology (CL) (Diehl et al., 2016) term. In cases where anatomical entities are defined primarily by their constituent cell types, we also provided the corresponding Uberon terms (e. g. mapping HPA “endothelial cell” to the Uberon term “Endotelium”). Cell clusters were mapped via their associated cell-type annotations. Overall, 81 out of 85 HPA cell types were successfully mapped to CL and/or Uberon terms. The unmapped annotations were “excitatory neuron”, “mixed cell types”, “mixed immune cell” and “undifferentiated cell”. As TS had a CL name annotation available, all 177 cell-types were successfully mapped to CL identifiers.

We compiled all known associations between HPO and CL terms, and between HPO and Uberon terms, with CoMent P-values < 0.001. Associations with P-values >= 0.001 (denoted as ‘-’ in the results) and HPO terms never co-mentioned with any cell type (denoted as “NA”) were considered negatives. Using this set as a gold-standard, we generatedReceiving Operator Characteristic (ROC) curves and computed the Area Under the Curve (AUC) to evaluate the performance of our method under all three criteria described previously.

## Code availability

All the code generated in this work is available at https://github.com/JNovoaR/Cell4Rare

