## Supplemental Figures for "Discovering cell types underlying rare disease phenotypes using scRNA-seq data from non-diseased tissues"

**Criteria for the manual annotation of phenotypes (HPO terms) to HPA or TS tissues**

To perform the manual mapping (b), we aimed to identify one or more broad HPO terms referring to abnormalities in the morphology and/or physiology of each tissue. We followed these criteria:

1. **Direct tissue-specific terms:** If available, we selected HPO terms explicitly referring to abnormalities of the given tissue (or one of its exact synonyms) using the format "Abnormality of the {tissue}," including both morphological and physiological abnormalities among their descendants (e.g., "Abnormality of the breast" for the tissues "Breast" [HPA] and "Mammary" [TS]).

2. **Separate morphological and physiological terms:** If a combined term was not available, we identified a pair of HPO terms separately covering morphological and physiological abnormalities of the tissue (or its synonyms).

3. **Broader anatomical terms:** If tissue-specific terms were unavailable, we selected HPO terms describing morphological and physiological abnormalities of broader anatomical entities that fully encompass the tissue.

4. **Morphological plus related physiological term:** If no broader physiological term was available, we selected a morphological HPO term as described above, together with a physiological term describing a biological function directly or partly performed by the tissue.

5. **Morphological term only:** If no clear physiological function could be ascribed to the tissue, we annotated it with the morphological HPO term alone (e.g., "Abnormality of the thymus").

6. **Medical test results:** When available, we added HPO terms referring to abnormalities detectable in medical tests performed on the tissue (e.g., cardiac or pulmonary tests).

Finally, we expanded the manual annotation by including all descendant terms of the selected HPO terms. A preliminary annotation was constructed by combining this manual mapping with the automated HPO-based annotation.

**TABULA SAPIENS SUPPLEMENTARY FIGURES:**

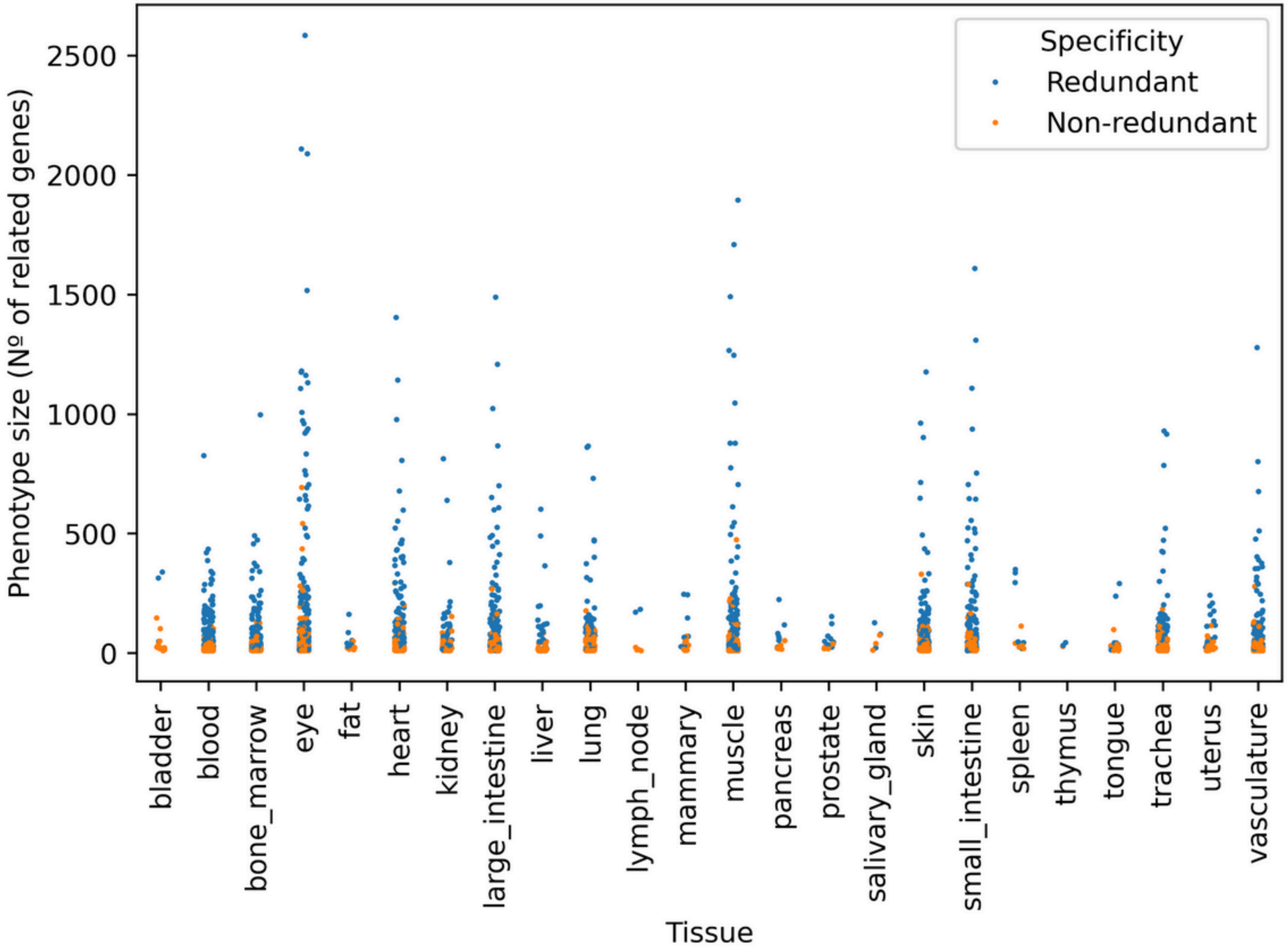


| Tissue | All | Non redundant |
|---|---|---|
| bladder | 14 | 11 |
| blood | 384 | 187 |
| bone_marrow | 230 | 119 |
| eye | 378 | 200 |
| fat | 14 | 7 |
| heart | 300 | 157 |
| kidney | 90 | 45 |
| large_intestine | 233 | 110 |
| liver | 55 | 28 |
| lung | 166 | 90 |
| lymph_node | 6 | 4 |
| mammary | 16 | 9 |
| muscle | 273 | 149 |
| pancreas | 17 | 8 |
| prostate | 16 | 6 |
| salivary_gland | 6 | 3 |
| skin | 284 | 156 |
| small_intestine | 235 | 112 |
| spleen | 11 | 5 |
| thymus | 3 | 1 |
| tongue | 22 | 15 |
| trachea | 160 | 88 |
| uterus | 34 | 19 |
| vasculature | 204 | 104 |
| Total tissue-phenotype pairs | 3151 | 1633 |
| Total unique phenotypes | 2193 | 1154 |

**Supplementary Figure S1**. **Distribution of phenotype sizes (left) and number of phenotypes per tissue (right) in the Tabula Sapiens (TS) dataset**. Phenotype size is defined as the number of HPO-associated genes expressed in at least one cell of the mapped tissue. Phenotypes are colored by specificity: **non-redundant** phenotypes have no child terms among those mapped to the same tissue, while **redundant** phenotypes have at least one child term. Some phenotypes are mapped to multiple tissues; the table shows both the total number of phenotype–tissue pairs (allowing repeats) and the total number of unique phenotypes.

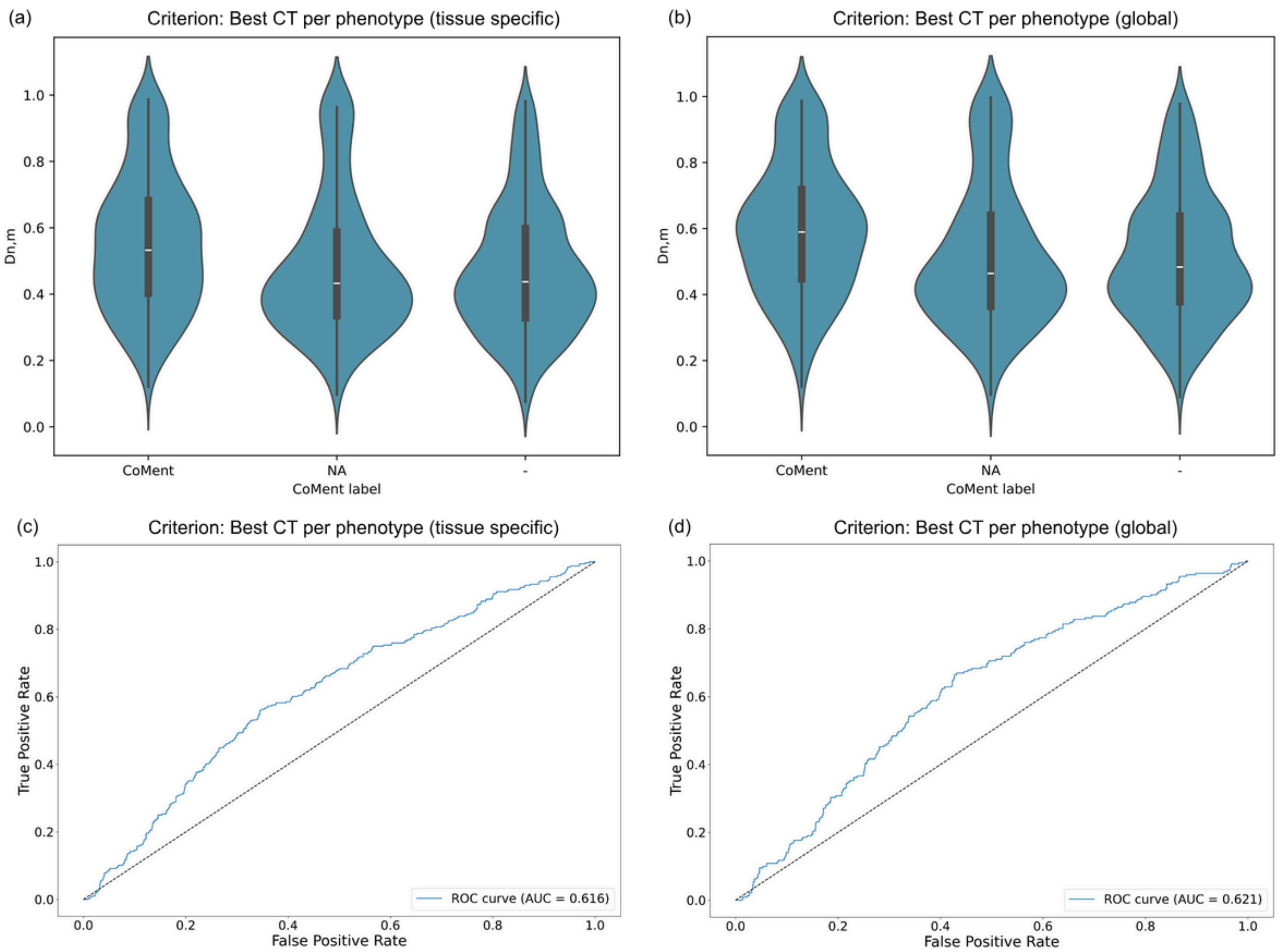


**Supplementary Figure S2. CoMent literature validation of the results of Cell4Rare applied over TS dataset**. Positive results were selected using two criteria: **(a, c)** the best cell type per phenotype–tissue pair (allowing multiple positives for phenotypes mapped to multiple tissues), and **(b, d)** the best cell type per phenotype (single positive per phenotype). Selected results were compared to phenotype–cell type pairs frequently co-mentioned in PubMed abstracts: **"CoMent"** indicates significant co-mention, **"–"** indicates non-significant or absent co-mention, and **"NA"** indicates no available information. Pairs tend to have higher $D_{n,m}$ values, especially under the global criterion. ROC curves and AUC values for both selection criteria are shown in (c) and (d). Performance was more modest than that observed using the Human Protein Atlas dataset.

**HUMAN PROTEIN ATLAS SUPPLEMENTARY FIGURES:**

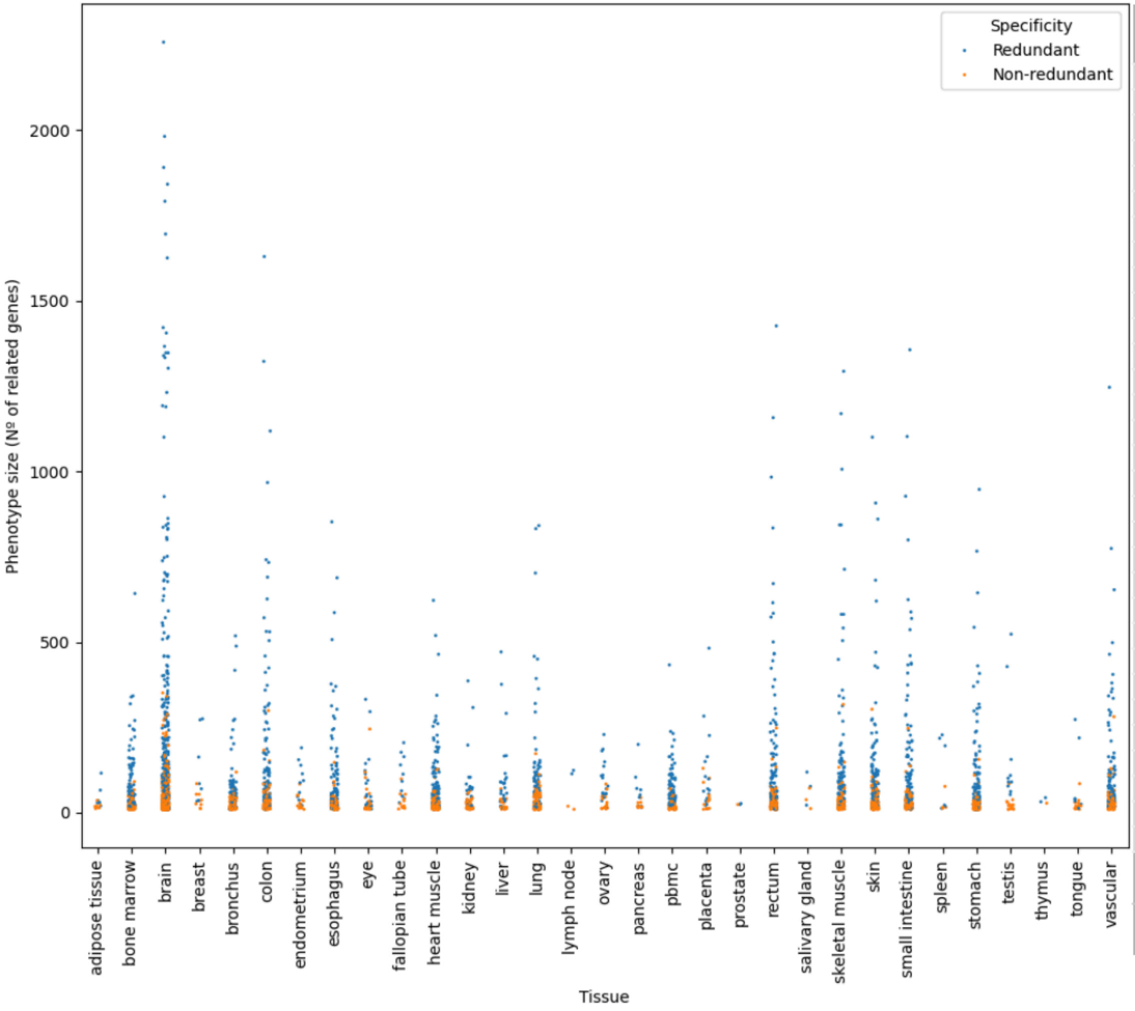


| Tissue | All | Non redundant |
|---|---|---|
| adipose tissue | 14 | 7 |
| bone marrow | 201 | 97 |
| brain | 872 | 462 |
| breast | 16 | 9 |
| bronchus | 133 | 71 |
| colon | 234 | 109 |
| endometrium | 24 | 13 |
| esophagus | 196 | 85 |
| eye | 59 | 35 |
| fallopian tube | 23 | 13 |
| heart muscle | 243 | 117 |
| kidney | 65 | 35 |
| liver | 45 | 21 |
| lung | 166 | 90 |
| lymph node | 4 | 2 |
| ovary | 33 | 15 |
| pancreas | 17 | 8 |
| pbmc | 166 | 79 |
| placenta | 30 | 14 |
| prostate | 16 | 6 |
| rectum | 232 | 109 |
| salivary gland | 6 | 3 |
| skeletal muscle | 254 | 137 |
| skin | 296 | 161 |
| small intestine | 224 | 103 |
| spleen | 10 | 4 |
| stomach | 204 | 89 |
| testis | 25 | 15 |
| thymus | 3 | 1 |
| tongue | 22 | 15 |
| vascular | 202 | 100 |
| Total tissue-phenotype pairs | 4035 | 1944 |
| Total unique phenotypes | 2503 | 1328 |

**Supplementary Figure S3. Distribution of phenotype sizes (left) and number of phenotypes per tissue (right) in the Human Protein Atlas (HPA) dataset**. Phenotype size is defined as the number of HPO-associated genes expressed in at least one cell of the mapped tissue. Phenotypes are colored by specificity: **non-redundant** phenotypes have no child terms among those mapped to the same tissue, while **redundant** phenotypes have at least one child term. Some phenotypes are mapped to multiple tissues; the table shows both the total number of phenotype–tissue pairs (allowing repeats) and the total number of unique phenotypes.

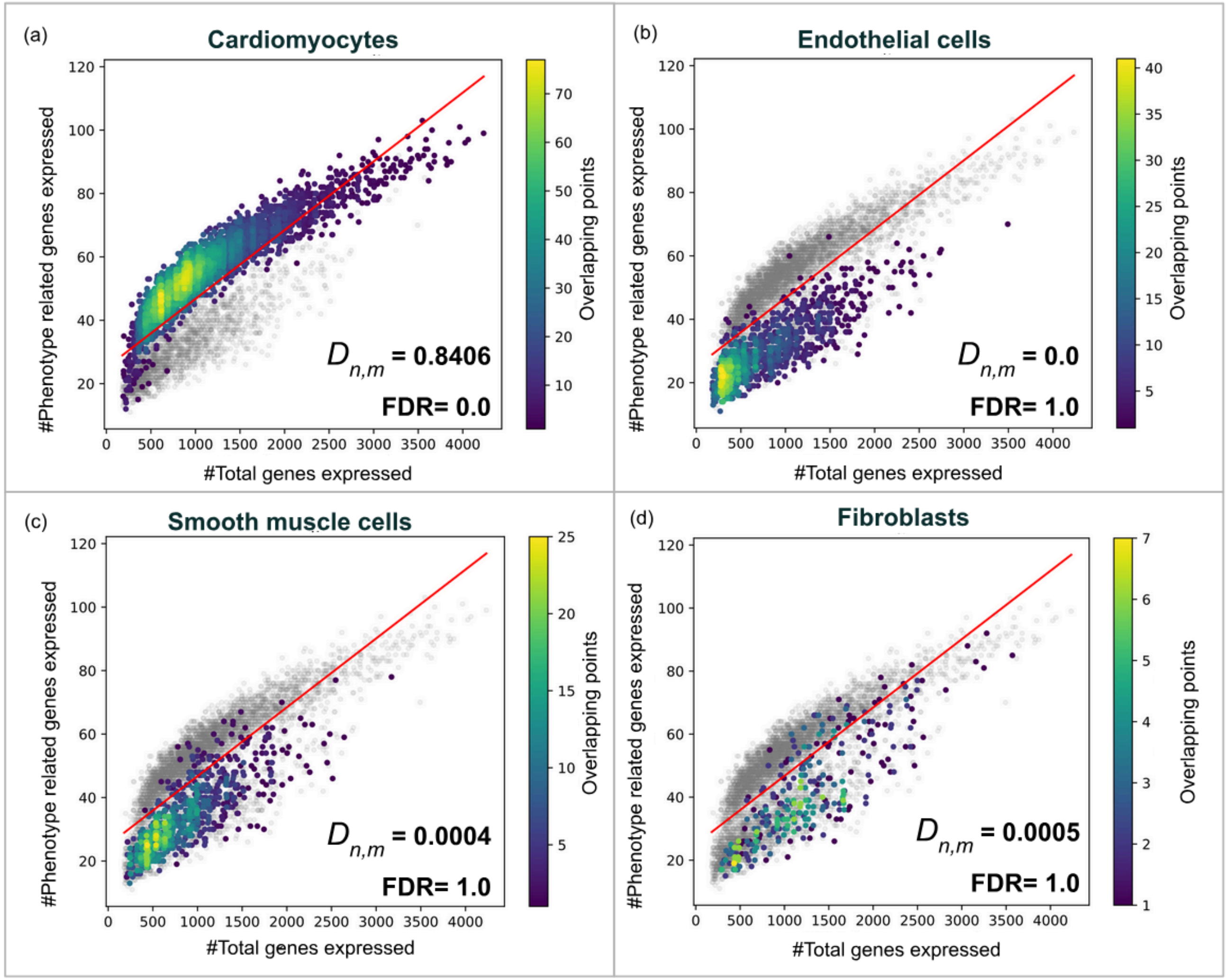


**Supplementary Figure S4. Cell4Rare results for dilated cardiomyopathy using HPA heart muscle scRNA-seq data**. **(a)** Cardiomyocytes show a significant association with dilated cardiomyopathy (FDR < 0.001), whereas **(b–d)** endothelial cells, smooth muscle cells, and fibroblasts show no significant association. Dilated cardiomyopathy is characterized by ventricular dilation and systolic dysfunction, often progressing to heart failure, and arises from diverse genetic etiologies, many of which impair cardiomyocyte structure and contractility.

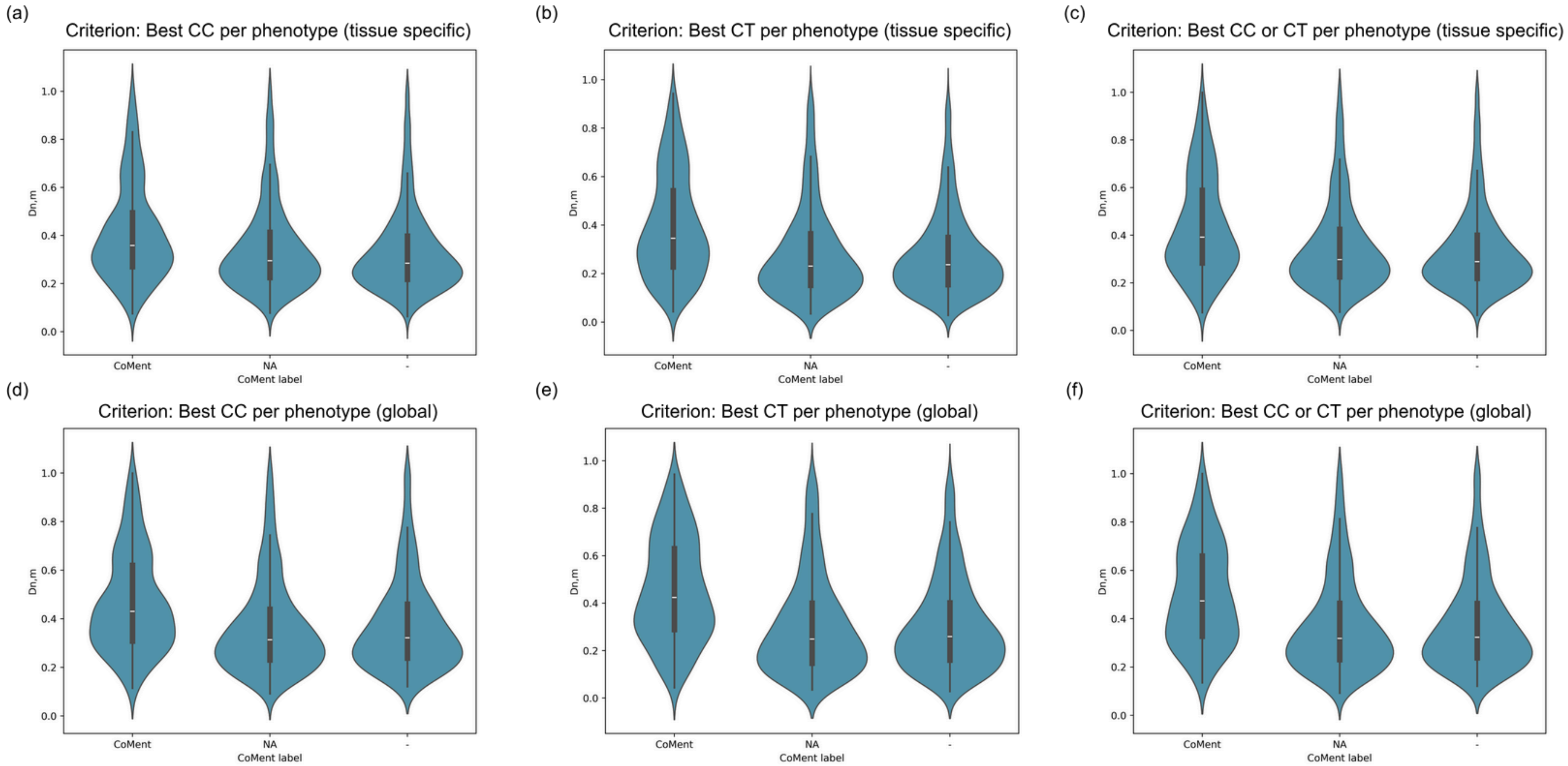


**Supplementary Figure S5**. **Distribution of Cell4Rare $D_{n,m}$ values (HPA dataset) for phenotype–cell type pairs supported or not supported by CoMent**. The gold standard consists of phenotype–cell type pairs frequently co-mentioned in PubMed abstracts: **"CoMent"** indicates significant co-mention, **"–"** indicates non-significant or absent co-mention, and **"NA"** indicates no available information. For Cell4Rare results, only the highest-scoring $D_{n,m}$ cell-type association per phenotype was selected. Selection involves two considerations: (1) choosing the top cell type, the cell type containing the top cluster, or the best between both; and (2) for phenotypes mapped to multiple tissues, selecting the best per tissue or the overall best across tissues. ROC analyses were performed across six combinations: **(a)** Best cluster per phenotype (tissue-specific), **(b)** Best cell type per phenotype (tissue-specific), **(c)** Best cluster or cell type per phenotype (tissue-specific), **(d)** Best cluster per phenotype (global), **(e)** Best cell type per phenotype (global), and **(f)** Best cluster or cell type per phenotype (global). Higher $D_{n,m}$ values are observed for pairs supported by CoMent, particularly under global criteria.

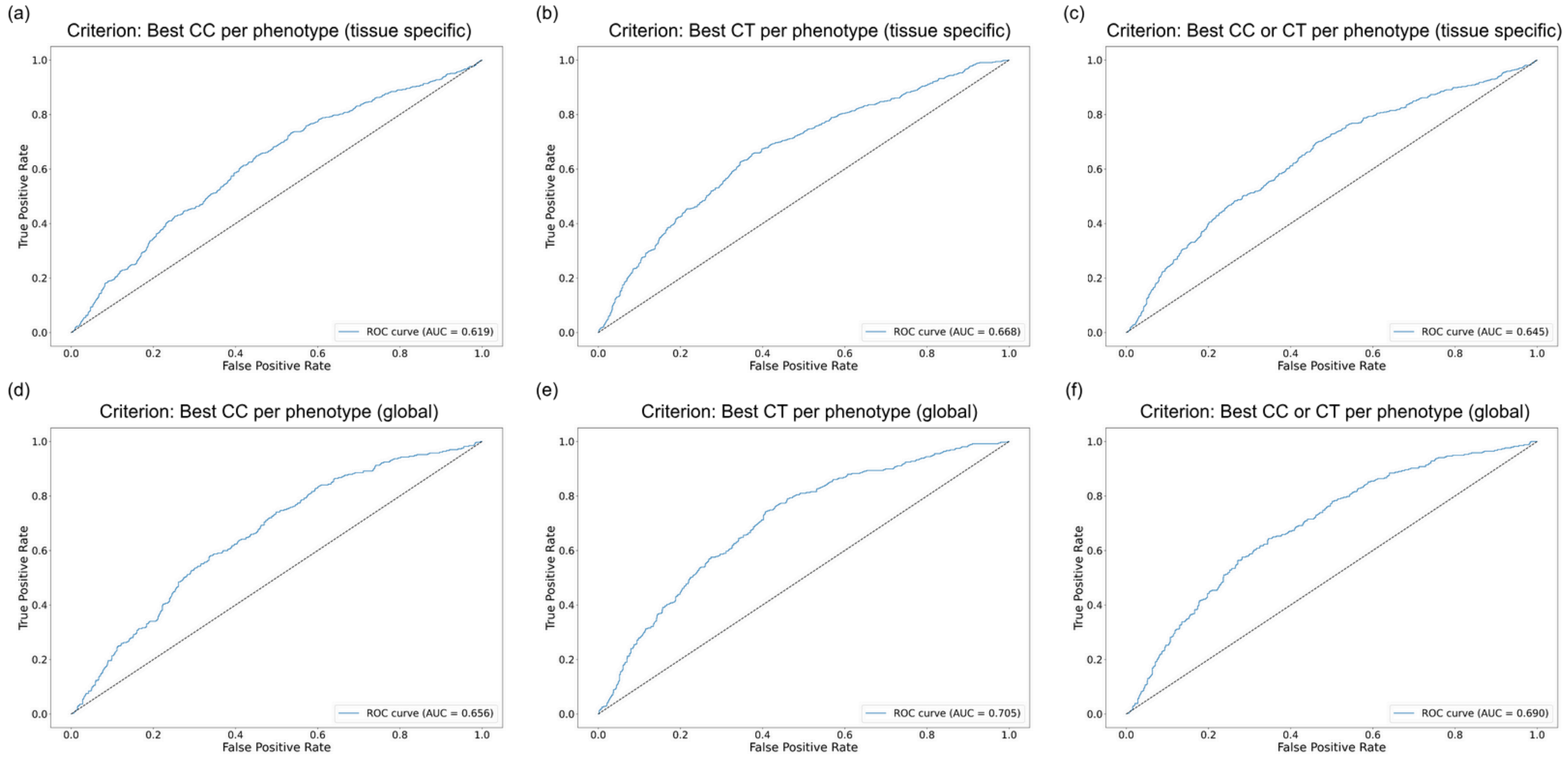


**Supplementary Figure S6**. **ROC curves and AUC values for CoMent literature validation of Cell4Rare results on the HPA dataset**. Positive results were defined as the highest-scoring ($D_{n,m}$) cell-type association for each phenotype. Selecting the best-scoring association involves two considerations: (1) choosing the top cell type, the cell type containing the top-scoring cluster, or the best between both; and (2) for phenotypes mapped to multiple tissues, selecting the best per tissue or the overall best across tissues. ROC analyses were performed across six combinations of these criteria: **(a)** Best cluster per phenotype (tissue-specific), **(b)** Best cell type per phenotype (tissue-specific), **(c)** Best cluster or cell type per phenotype (tissue-specific), **(d)** Best cluster per phenotype (global), **(e)** Best cell type per phenotype (global), and **(f)** Best cluster or cell type per phenotype (global). Global and Best Cell-Type criteria consistently performed best, with peak performance achieved using the Best Cell-Type per phenotype (global) criterion (AUC = 0.705).

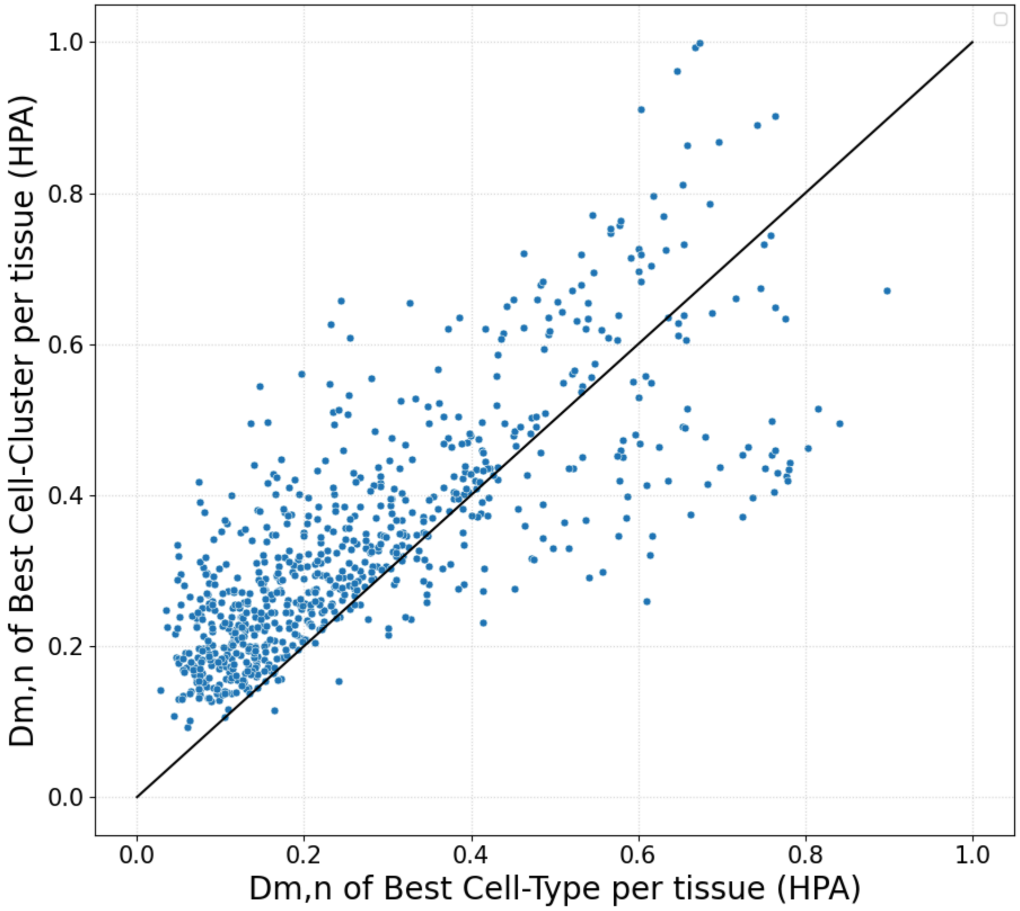


**Supplementary Figure S7. Comparison of $D_{n,m}$ values for the best cell-type versus the best cell cluster per phenotype in HPA.** Each point represents a phenotype mapped to an HPA tissue, with x as the $D_{n,m}$ value of the best cell type and y as that of the best cell cluster. Black line: x=y. Phenotypes where the best cell type and the best cell cluster are identical (i.e., the cell type consists of a single cluster) were excluded for clarity.

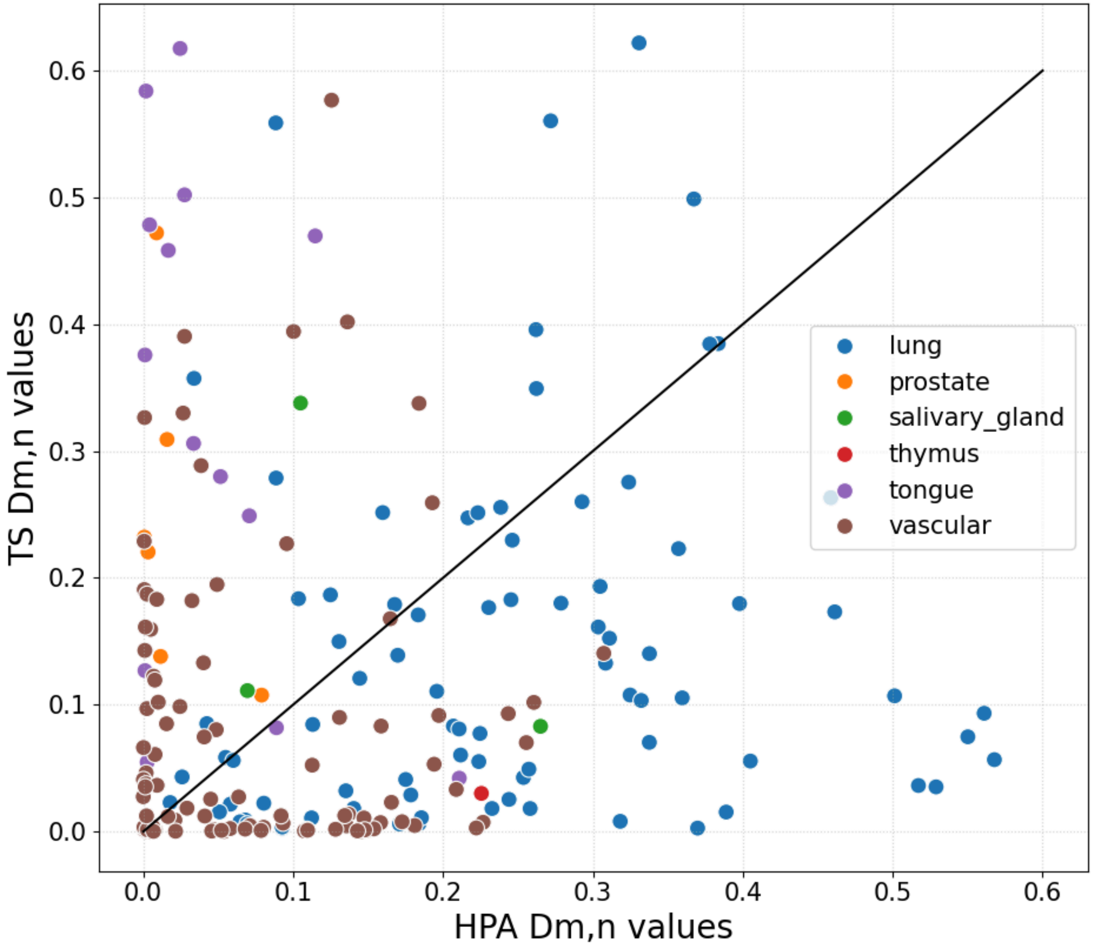


**Supplementary Figure S8. Impact of cell-type annotation on Cell4Rare results**. Comparison of $D_{n,m}$ values for the best cell type per non-redundant phenotype between HPA and TS for the six HPA tissues derived from TS. The underlying scRNA-seq data are identical, but cell annotations differ. Results show poor correlation in most cases, highlighting the influence of cell-type annotation on Cell4Rare outcomes. **Black line:** x=y.
